# Epoxyazadiradione ameliorates Parkinson’s disease by upregulating heat shock factor 1 and protein degradation pathways in mice

**DOI:** 10.64898/2025.12.08.692835

**Authors:** Shivani Chandel, Naibedya Dutta, Hossainoor Rahaman Sareng, Nilanjana Ghosh, Izaz Monir Kamal, Anirban Manna, V. Ravichandiran, Jayanta Mukhopadhyay, Saikat Chkrabarti, Rajiv Kumar Goswami, Mahadeb Pal

## Abstract

Parkinson’s disease (PD) is a major debilitating health concern for millions of the elderly population all over the world. This progressive neurodegenerative disorder also poses a severe mental and financial burden to caregivers and society. Despite a major thrust on research for therapy development, no significant progress has been made; only temporary management options are currently available. To this end, we have reported azadiradione (AZD), a triterpenoid that we isolated from neem seed extract using a cell-based assay. AZD showed high efficacy in ameliorating protein aggregation-induced pathology and symptoms in fruit flies and mice. Current evidence suggests that AZD functions through activating the transcriptional function of heat shock factor 1 (HSF1), a master regulator of protein quality control pathways, without modulating the cellular redox balance. To better understand the pharmacophore of AZD, a triterpenoid in its observed function, we have analysed various structural derivatives, focusing on their HSF1-activating function in vitro and their efficacies in ameliorating protein aggregation-induced toxicities in cell and mouse models. Our analyses, based on real-time PCR, immunoblots, fluorescent anisotropy, and a mouse model of MPTP-induced PD, highlighted Epoxy-azadiradione (Epoxy) as being as efficient as AZD in in vivo functional tests, albeit activating the promoter binding activity of HSF1 with at least two-fold higher efficacy in vitro. Notably, similar to AZD, Epoxy did not induce cellular redox imbalance. We also incorporated molecular docking analyses involving the published crystal structure of the DNA-binding domain of HSF1 bound to its DNA recognition element to study molecular dynamics-based energy estimation. The analysis revealed a higher energy stability of the epoxy-bound complexes, as indicated by a significant decrease in binding free energy (ΔG) estimated from an ensemble of intermediate docked complex structures.

## 1 Introduction

Parkinson’s disease (PD), the second most common neurodegenerative disorder after Alzheimer’s disease, currently affects about 6 million people worldwide (Dorsey et al., 2018; Reeve et al., 2014). Data indicate that the elderly population, generally those 65 years and above, becomes more vulnerable to the disease as they age (Dorsey et al., 2018; Reeve et al., 2014). About 5% of the population included in the susceptible categories were reported to show PD symptoms even before the age of 60 (Schulte and Gasser et al., 2011). Clinical features of PD include bradykinesia, rigidity, tremor, gait dysfunction, and postural instability (Armstrong and Okun et al., 2020; Klein and Westenberger et al., 2012). A loss of approximately 60% of dopaminergic neurons (DA) in the substantia nigra pars compacta, and a decrease of up to 70-80% of normal DA content in the striatum, were observed in the clinical stage of PD pathology (Larkov et al., 2020). PD occurs in familial or sporadic forms; about 10% of cases are familial/hereditary, while the rest are sporadic or idiopathic with a largely unknown cause (Klein and Westenberger et al., 2012). Hereditary PD has been linked to at least one defect in several genes. A mutation in the SNCA/PARK1-4 or LRRK2/PARK8 gene is associated with the autosomal dominant form, while mutations in Parkin/PARK2, PINK1/PARK6, DJ-1/PARK7, or ATP13A2/PARK9 are associated with the recessive form of the disease. Sporadic PD has been linked to at least the mutation in the glucocerebrosidase (GBA) gene, the SNCA REP1 promoter polymorphism, or the G2385R and S1647T variants in the LRRK2 gene (Kluss et al. 2019; Lunde et al. 2018; Kluss et al. 2019; Selvaraj and Piramanayagam 2019). A mutation in the gene encoding α-synuclein has been reported as a major common pathogenic factor in PD. A major pathological signature of PD is the deposition of insoluble α-synuclein aggregates called Lewy bodies (LB) as nuclear inclusions. In addition to α-synuclein, the LB contains molecular chaperones and components of protein degradation pathways, suggesting their involvement in protein quality control (PQC) pathways in maintaining protein homeostasis in normal/healthy cells (Balchin et al., 2016; Mehra et al., 2019).

PQC, through the coordinated activation of molecular chaperones and protein degradation pathways, protects cells from toxicity caused by the formation of misfolded proteins (Upadhyay et al., 2021). The molecular chaperones primarily comprise the inducible heat shock proteins (HSPs), which regulate protein quality by maintaining the native conformations of proteins through their guidance of folding, assembly, disassembly, transport, and degradation (Upadhyay et al., 2021). The protein degradation pathways, the ubiquitin-proteasome system (UPS) and the autophagy-lysosomal pathway (ALP), serve an indispensable function in PQC by clearing cellular protein aggregates through their enzymatic functions (Dikic et al., 2017). Preferential degradation of α-synuclein/protein aggregates by the UPS or ALP suggests highly regulated cross-talk between these pathways (Cuervo et al., 2004; De Mattos et al., 2020; Lee et al., 2004; Paxinou et al., 2001; Vogiatzi et al., 2008; Webb et al., 2003; Yu et al., 2009). While UPS degrades the small α-synuclein aggregates, the relatively large aggregates are targeted for degradation by the ALPs (Ebrahimi-Fakhari et al. 2011). UPS mediates the selective degradation of different α-synuclein aggregates through an ATP-dependent, multi-subunit 26S proteasome, as demonstrated by both in vitro and in vivo studies (De Mattos et al., 2020). The catabolic process of ALP converges at the lysosome. The major form of autophagy, macroautophagy, involves the sequestration of substrates in an autophagosome, a double-layered membrane-bound structure, to target lysosomal degradation (Sontag et al., 2017). Defects in genes such as ATG5 in the ALP cause neurological dysfunction. Overexpression of beclin 1, another gene in the ALP, can rescue α-synuclein-induced neurological defects in mice (De Mattos et al., 2020). Evidence supports the idea that the HSP70 molecular chaperone modulates activity of both UPS and ALP pathways; HSP70, through its co-chaperone carboxyl terminus of Hsp70-interacting protein (CHIP), a E3-ubiquitine ligase, facilitates the degradation of α-synuclein by both UPS and ALP (Kampinga and Craig et al., 2010; Shin et al., 2005).

Heat shock factor 1 (HSF1), a transcription activator for the inducible heat shock protein (Hsp) genes such as Hsp70 and Hsp90, is considered a master regulator of the heat shock response. In a normal/unstressed cell, monomeric HSF1 is sequestered in a repressive complex in the cytoplasm in association with Hsp90, Hsp70, Hsp40, p23, J-proteins, and TRiC/CCT (Feder et al., 2021; Zheng et al., 2016). Exposure of a cell to a thermal shock or other proteotoxic stress results in the dissociation of this repressive complex due to titration away of the chaperones by the misfolded proteins; the monomeric HSF1 trimerizes and translocates to the nucleus to bind to its recognition sequence, heat shock elements (HSE) constituted with three to eight units of 5’-nGAAn-3’ on the promoters of its target genes (Morimoto et al., 1998; Zheng et al., 2016; Ali et al., 2019). Posttranslational modifications (PTMs), such as phosphorylation, acetylation, and sumoylation, are important for the activation of HSF1 functions (Anckar et al., 2011). However, HSF1 lacking a regulatory domain has been found to respond well to stress with a reduced activation threshold. HSF1 consists of an amino-terminal winged domain that activates and inhibits HSF1; site-specific acetylation modulates HSF1 protein stability and/or DNA-binding property. Under basal conditions, HSF1 is stabilized by p300-induced acetylation at Lys208 and Lys298, which prevents proteasomal degradation (Gomez-Pastor et al., 2018). p300-mediated Lys80 acetylation inhibited HSF1 from directly interacting with HSE, a process counteracted by SIRT1, a NAD-dependent protein deacetylase (Gomez-Pastor et al., 2018). HSF1 was shown to undergo multiple phosphorylation events on different residues, including those mediated by p38 MAPK, which phosphorylates HSF1 at Ser326 (Kmiecik et al., 2022).

Interestingly, different protein kinases can lead to phosphorylation of the same residues in various contexts. In Huntington’s disease, phosphorylation at Ser303 and Ser307 by casein kinase 2 (CK2) leads to the inactivation and degradation of HSF1, mediated by ubiquitylation in neurons (Thiele et al., 2018). Sumoylation of HSF1 protein at Lys298 residue led to a compromised HSF1 activation and decreased hsp70 induction (Thiele et al., 2018). Phosphorylation at Ser326 by various kinases, including DNA-PK, ERK1/2, mTOR, and PI3K, is implicated in HSF1 activation (Tye and Churchman 2021; Simoncik et al. 2024).

Because the accumulation of protein aggregates, including α-synuclein, has been attributed to the pathology of PD, several laboratories have demonstrated that forced upregulation of HSF1 or its target Hsp70 ameliorated disease symptoms and associated pathology (Nelson V et al., 2016; Singh B et al., 2018). We have previously identified azadiradione (AZD, 451.5 Da) as a unique activator of HSF1, which ameliorates protein aggregation-induced toxicity in fruit flies, Huntington’s disease in mice (Singh et al. 2018) , as well as MPTP-induced PD in mice with significant efficacy(Sareng et al. 2025). AZD interacts physically with HSF1 to enhance its binding to the HSE in vitro (Nelson V. et al., 2016). To improve the functional effectiveness of the triterpenoid (AZD) in vitro and in vivo, we investigated the role of its functional groups (pharmacophores) in its observed activity. Through analysing a group of structural derivatives, we identified an epoxy derivative of AZD, called Epoxy-azadiradione (Epoxy), with more than two-fold greater functional efficacy in mitigating protein aggregation-associated toxicity.

## 2 Materials and methods

### 2.1 Isolation of azadiradione and its natural derivatives from Azadirachta indica

The flaked Azadirachta indica seeds were extracted with methanol for 4 days at room temperature. The methanolic extracts were combined and concentrated using a rotary evaporator at 50 °C to yield a brownish oil. The methanol extract was re-extracted with hexane, dichloromethane, and ethyl acetate, and then evaporated to yield a brown gum. The resultant dichloromethane fraction was subjected to gross column chromatography with a hexane-ethyl acetate solvent system. The fractions eluted with hexane-ethyl acetate (80:20 and 70:30) solvent systems were pooled according to their TLC profiles and concentrated to yield a fraction, which was then subjected to silica gel column chromatography using a 5% isopropanol in hexane eluent to isolate the compounds. The purity of the compounds was monitored by HPLC. The identification of the compounds was confirmed by recording the ¹H and ¹³C NMR spectra at 500 and 125 MHz, respectively, on a Bruker DRX 500 spectrometer using CDCl₃ as the solvent. The δ values are reported in ppm (TMS as the internal standard), and the J (Hz) assignments of the ¹H resonance coupling are provided.

### 2.2 Synthesis of azadiradione derivatives

1. 2 mL of an acetone solution of AZD (10 mg, 22 µM) was stirred at 0°C with 50 ml of 5% solution of osmium tetroxide in n-butanol (1 µmol, 0.05 molar equivalent) and a 1 M solution of N-methyl morpholine N-oxide (3.86 mg, 33 µmol, 1.5 molar equivalents) for overnight. The reaction was quenched by stirring with a saturated sodium sulphite solution for 1 h at room temperature. The reaction mixture was then extracted with ethyl acetate three times. The combined organic layers were dried to obtain the crude product, which was then subjected to preparative TLC using a hexane: ethyl acetate (60:40) solvent system to isolate compound S2 (21%).
2. A 2 mL DCM solution of S2 (10 mg, 20 µM, one molar equivalent) was stirred at room temperature with 2,2-dimethoxypropane (10 µL, 88 µmol, 4.4 molar equivalents), followed by the addition of catalytic camphorsulfonic acid. A saturated solution of sodium bicarbonate quenched the reaction. The reaction mixture was extracted 3 times using DCM. The combined organic layers concentrated by vacuum evaporation were subjected to preparative TLC using a hexane: ethyl acetate (80:20) solvent system to obtain compound S3 (32%) m/z: 524.64, C31H40O7.
3. To a methanolic solution of AZD (10 mg dissolved in 1 ml methanol, 22 µM) at 0°C under nitrogen, was added finely powdered potassium carbonate (3.06 mg, 22 µM). The reaction mixture was stirred at room temperature for 12 h. The completion of the reaction was tested by monitoring the products by TLC. The crude product, concentrated by vacuum evaporation of the solvent, was subjected to preparative TLC using a hexane: ethyl acetate (60:40) solvent system to obtain compound S4 (50%). m/z: 408.52, C26H32O4.
4. AZD (in methanol, 10 mg, 22 µM) was stirred at room temperature under nitrogen with six molar equivalents of sodium tetraborohydrate (5 mg, 132 µM) for 3 h. The reaction after quenching with a saturated ammonium chloride solution was extracted three times with ethyl acetate. The combined organic layers were dried, and the crude product was isolated by preparative TLC using ethyl acetate (65:35) to obtain compound S6 (60%), m/z: 458.62, C28H42O5.
5. AZD (10 mg, 22 µM in ethanol) was added to a solution containing sodium acetate (3.06 mg, 374 µmol, 1.7 molar equivalent) and hydroxyl ammonium chloride (3.82 mg, 55 µmol, 2.5 molar equivalent) at room temperature, and the resultant mixture was refluxed for 1 day. The completion of the reaction was confirmed by TLC monitoring of the products. The crude product collected by vacuum evaporation was subjected to preparative TLC using a hexane: ethyl acetate (60:40) solvent system to isolate compound S7. m/z: 465.58, C28H36N2O5.
6. AZD (10 mg, 22 µM) was added with 5% Pd/C (1.2 mg) in 1 ml ethyl acetate with repeated stirring under hydrogen at room temperature. The fully purged flask was stirred at room temperature for 12 h. After the reaction was completed, the product was passed through a Celite column, and the excess solvent was evaporated. The crude product was purified by preparative TLC using hexane: ethyl acetate (80:30) solvent system to isolate compound S8 m/z: 456.61, C28H40O5.
7. AZD (10 mg, 22 µM in acetic anhydride), catalyst p-toluene sulphonic acid, was added at room temperature, and the resultant mixture was stirred overnight. The completion of the reaction was confirmed by TLC monitoring of the products. The crude product collected by vacuum evaporation was subjected to preparative TLC using a hexane: ethyl acetate (60:40) solvent system to isolate compound S9. m/z: 481.58, C29H36O6.

### 2.3 Cell culture

HEK 293 (Human embryonic kidney) and Neuro-2A (Mouse neuroblast) cells were cultured in DMEM supplemented with 10% fetal bovine serum, penicillin (50 μg/ml), streptomycin (50 μg/ml), and non-essential amino acids and maintained in a humidified incubator at 37 °C (Dutta et al., 2021).

### 2.4 Cell viability assay

Cell viability was assessed using the MTT (3-(4,5-dimethylthiazol-2-yl)-2,5-diphenyltetrazolium bromide) assay according to the standard protocol. Cells seeded in 96-well plates were treated with different concentrations of compounds or vehicles for 24 h. Next, the growth media was replaced with MTT at 0.5 mg/mL and incubated at 37°C for 4 h. The MTT solution was then discarded, and the purple-colored formazan crystals were dissolved in DMSO to measure the absorbance at 570 nm using a microplate reader.

### 2.5 RNA isolation and quantitative PCR

Total RNA was isolated from cells using TRIzol according to the manufacturer’s protocol (Invitrogen). RNA was estimated using a Nanodrop spectrophotometer. One microgram of total RNA was transcribed using the iScript cDNA synthesis kit (Bio-Rad). RT-qPCR was performed using SYBR green (Applied Biosystem) (Dutta et al., 2021).

### 2.6 Flow cytometric analysis of intracellular ROS

The ROS production was evaluated using 2,7-dichlorofluorescein diacetate (DCFDA) fluorescence dye. The cells were treated with varied concentrations of compounds overnight. The cells were treated with 5 μM DCFDA (Merck) at 37 °C for 30 minutes in the dark. All the cells were then harvested and subjected to flow cytometry (Dutta et al., 2021).

### 2.7 Cell lysate preparation and western blotting

Mouse neuroblastoma Neuro-2A cells were harvested and pelleted after washing with ice-cold PBS after the desired treatments. The cells were lysed with lysis buffer containing 20 mM Tris-HCl (pH 7.5), 1% Triton X-100, 150 mM NaCl, 5% glycerol, 1 mM phenylmethyl sulfonyl fluoride (PMSF), 10 μg/ml leupeptin, 10 μg/ml aprotinin, 20 mM sodium fluoride, and 20 mM sodium orthovanadate. Protein estimation of the whole cell lysate was performed using the Bradford assay reagent (Bio-Rad) with BSA as a standard. The proteins in the whole-cell lysates (15 µg) were boiled in Laemmli buffer after resolving by SDS-PAGE and then transferred onto a PVDF membrane. This was followed by blocking with 5% milk in PBST (PBS plus 0.5% Tween 20). The membranes were incubated overnight at 4 °C with an appropriate dilution of primary antibody (prepared in PBST supplemented with 5% BSA). They were then washed with PBST and incubated with a secondary antibody in PBST supplemented with 5% BSA. After 1 h of incubation at room temperature, the membrane was washed with PBST, followed by incubation with ECL to develop under a ChemiDoc (Bio-Rad) (Dutta et al., 2021).

### 2.8 Fluorescence polarization assay

Human HSF1 protein purified from an overexpressing E. coli strain, attached with a hexa-histidine tag at its N-terminus, and fluorescently labelled double-stranded oligonucleotides as Heat shock element (HSE) (with 10 nM 5 5’-fluorescein amide modification) were used for fluorescence polarisation (FP) studies. FP experiments were performed in FP buffer (25 mM HEPES, pH 7.4, and 75 mM NaCl), and the baseline millipolarization value was taken with (Excitation at 495 nm and emission at 517nm). AZD and its derivatives were titrated in the reaction with millipolarization values taken at a constant HSF1 concentration. Kd values were calculated by plotting the curves from three independent experiments using the one-site specific binding fit of the curves in GraphPad Prism 7 software.

### 2.9 In vivo experiment

Adult male Swiss albino mice (4-6 weeks old) weighing around 22-25 g were acclimated for a week in a controlled environment at 22°C, 50% humidity, under 12-hour light/dark conditions with ad libitum access to food and water. Mice were randomly divided into six groups, each containing 5 animals (n = 5). MPTP was used to develop the PD model due to its ability to destroy dopaminergic neurons by apoptosis in the substantia nigra pars compacta (Schildknecht et al. 2017). Before MPTP administration, the weight and grip strength of the mice were noted. Groups A, B, and C were control groups treated with saline or drugs only: Group A consisted of control mice, Group B of Epoxy–treated mice, and Group C of AZD-treated mice. The PD model was developed in Groups D, E, and F mice by treating them with MPTP on alternate days for seven days (10 mg/kg, i.p.). The functional efficacy of compounds in the PD model was tested by treating Groups E and F with Epoxy and AZD (10 mg/kg), respectively, on alternate days for 20 days. During the total treatment period, the body weight and grip strength of each mouse were observed on alternate days (Dutta et al., 2021). The experiment was carried out with prior permission from the BI animal ethics committee (IAEC/BI/88/2018).

### 2.10 Immunofluorescence and immunohistochemistry

Neuro-2A cells were grown to 70% confluency on grease-free coverslips in a microplate. Cells were fixed with 4% paraformaldehyde in PBS by incubation for 20 min at room temperature, followed by three washes with ice-cold PBS. For permeabilization, the samples were incubated with 0.1% Triton X-100 in PBS for 10 min at room temperature, followed by three washes with PBS. Cells were treated with 3% BSA in PBS-Tween 20 for 30 min. After three washes with PBS (each with a 5-minute incubation), the samples were incubated with the secondary antibody for 2 hours in the dark. Cells were then counterstained with DAPI and mounted on coverslips for imaging under a confocal microscope (Leica). Mouse brain samples were embedded in paraffin blocks and sectioned by microtome. For deparaffinization, paraffin-embedded sections were washed with xylene and then processed sequentially with xylene: ethanol (1:1), 100% ethanol, 90% ethanol, 70% ethanol, and 50% ethanol. Immunohistochemistry with anti-tyrosine hydroxylase (TH) antibody was performed by using an IHC staining kit from Vector Laboratories (Dutta et al., 2021).

### 2.11 Molecular docking of Azadiradione and Epoxyazadiradione in the HSF1-HSE complex

Molecular docking analyses of AZD and Epoxy were performed using meta-docking programs AutoDock v4.2 (Morris et al., 2009), LeDOCK (Liu et al., 2019), and rDOCK (Ruiz-Carmona et al., 2014) (Figure S1). Best-ranked docked solutions (top 15) from all three programs were pooled together to score further using the AutoDock Vina (Trott et al., 2010) program. Docking solutions were clustered based on the positional similarity represented by the root mean square deviation (RMSD) of the docked ligands. Docked complexes having RMSD <= 2.5Å are grouped into a single cluster. Energetically preferred 3D conformers of AZD and Epoxy retrieved from PubChem ( Kim et al., 2016) were generated using the FROG2 (Free Online Drug conformation generation) software (Miteva et al., 2010) for docking in one of the crystal structures of HSF1-HSE complex available in the protein data bank (PDB) (PDB ID: 5D5U) (Neudegger et al., 2016). Initially, a blind/naïve docking approach was implemented to find out the most preferred/populated binding pockets within the HSF1-HSE monomer complex. Two distinct binding pockets/sites were found to be mostly populated. Site 1 is mostly comprised of a helix-loop-helix binding region, whereas the protein-DNA interface site constructs site 2 (Figure S1A). CHIMERA (Pettersen et al., 2004) and LIGPLOT (Wallace et al., 1995) programs were used to generate the protein-ligand interaction complex images. The highest rank docking solution from the most significant clusters was selected, and docking scores were compared (Figure S1B).

### 2.12 Molecular dynamics analyses

The 3D structures of HSF1-HSE docked with AZD and Epoxy were subjected to molecular dynamics (MD) simulation using the GROMACSv4.5.3 simulation package (Abraham et al., 2015) to understand the structural and energetic stabilities of the proposed protein-DNA-drug complexes. Coordinates and topology files of the receptor molecules were generated with the Amberff99sb force field (Wang et al., 2004). The topology and coordinate files of ligands were generated using ACPYPE (AnteChamber Python Parser interface) software (Sousa da Silva et al., 2012). Steepest-descent (Nocedal and Wright et al., 2006) and conjugate-gradient (Straeter et al., 1971) minimization algorithms were utilized for complexes embedded within a TIP3P water-filled simulation box (Mark and Nilsson et al., 2001). Equilibration was done under NVT (constant number of particles, volume, and temperature) and NPT (constant number of particles, pressure, and temperature) conditions. Trajectories were saved at an interval of 0.02 ps, and a total of 500,000 snapshots were recorded from a 10 ns simulation at a temperature of 300 K and a pressure of 1 atm.

### 2.13 Estimation of the binding free energy of docking complexes

The free energy of the HSF1-HSE/Epoxy docking complexes was estimated using a molecular mechanics approach combined with the Poisson-Boltzmann and surface area (MM-PBSA) method (Genheden et al., 2015). The gmx_MMPBSA tool was used to assess the binding affinity of the docking complexes derived from ensemble structures based on molecular dynamics simulations (Valdés-Tresanco et al. 2021). Binding free energies were calculated from 40 snapshots extracted at an interval of 250 ps from the 10 ns simulation trajectory. The Interaction Entropy (IE) module was used for calculating the entropic component of the binding free energy (Duan et al. 2016). Figure S1C provides an overview of the MD simulation, followed by the energy estimation protocol implemented for the HSF-HSE/Epoxy complexes.

## 3 Results

### 3.1 Synthesis and isolation of azadiradione derivatives (**Figure 1**)

We have isolated and characterized four natural compounds from neem seeds having structural similarity as Azadiradione [(5R,7R,8R,9R,10R,13S,17R)-17-(furan-3-yl)-4,4,8,10,13-pentamethyl-3,16-dioxo-6,7,9,11,12,17-hexahydro-5H-cyclopenta[a]phenanthren-7-yl] acetate, Epoxy-azadiradione [(1S,2R,4S,6S,7S,10R,11R,16R,18R)-6-(furan-3-yl)-1,7,11,15,15-pentamethyl-5,14-dioxo-3-oxapentacyclo[8.8.0.02,4.02,7.011,16]octadec-12-en-18-yl] acetate ], Gedunin [(1S,2R,4S,7S,8S,11R,12R,17R,19R)-7-(furan-3-yl)-1,8,12,16,16-pentamethyl-5,15-dioxo-3,6-dioxapentacyclo[9.8.0.02,4.02,8.012,17]nonadec-13-en-19-yl] acetate], Azadirone[(5R,7R,8R,9R,10R,13S,17R)-17-(furan-3-yl)-4,4,8,10,13-pentamethyl-3-oxo-5,6,7,9,11,12,16,17-octahydrocyclopenta[a]phenanthren-7-yl] acetate].

**FIGURE: 1.**
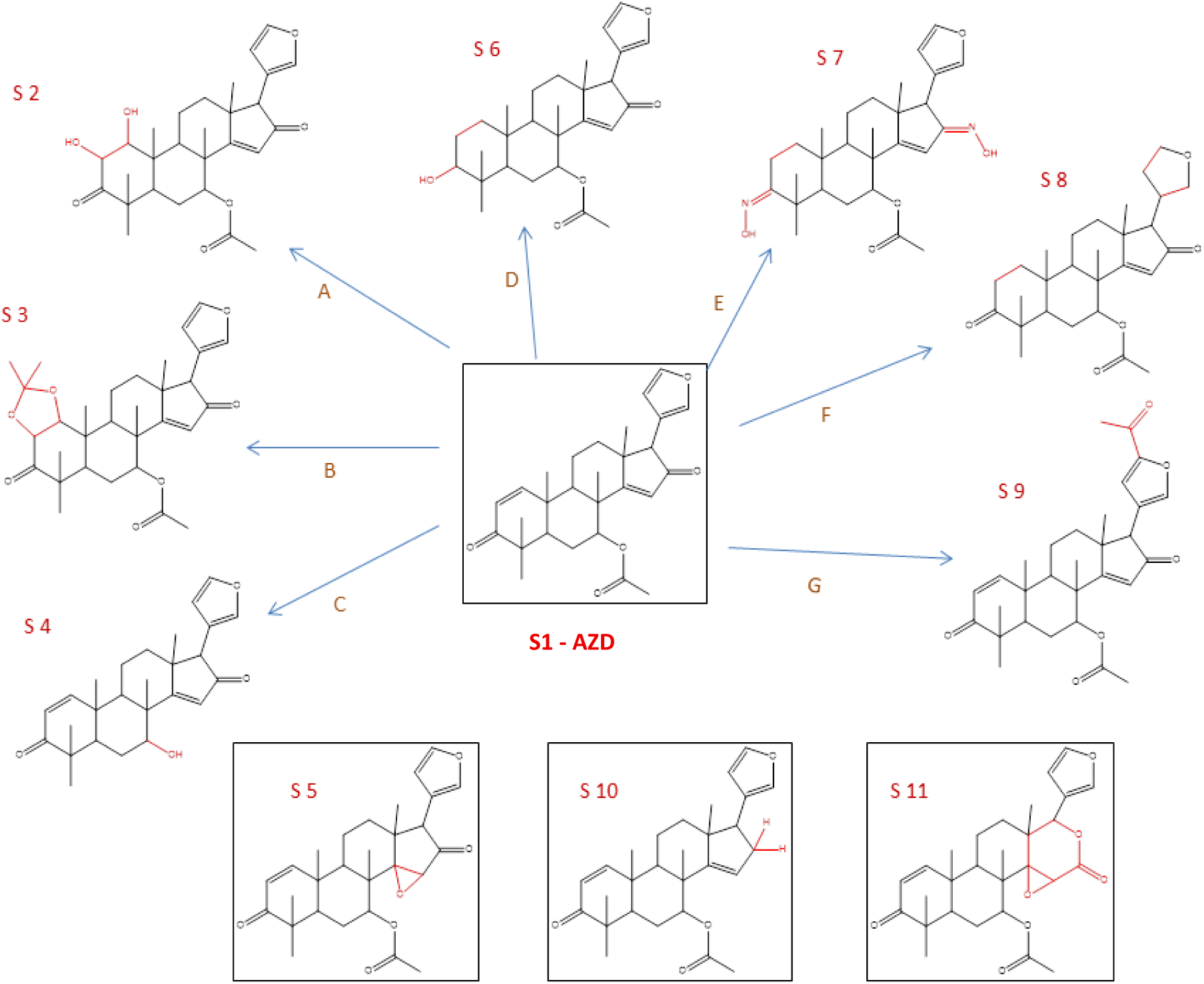
**Chemical structures of the derivatives of azadiradione synthesized or isolated from neem seed extracts.** Derivatives synthesized from neem seeds are boxed. Azadiradione is shown at the centre. The synthesis and identification methods are described in the Materials and Methods.

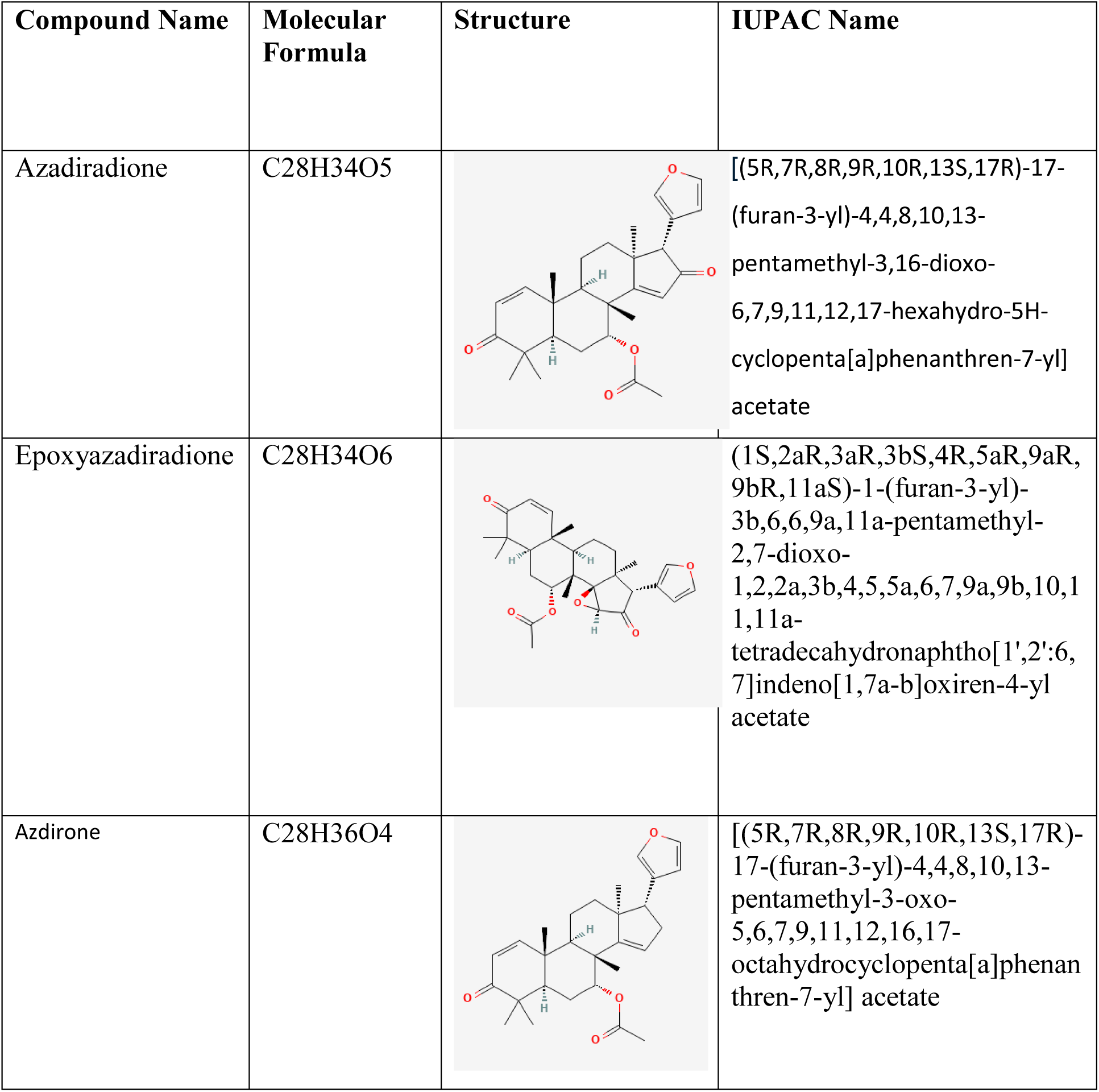

| Compound Name | Molecular Formula | Structure | IUPAC Name |
| --- | --- | --- | --- |
| Azadiradione | C<sub>28</sub>H<sub>34</sub>O<sub>5</sub> |  | [(5R,7R,8R,9R,10R,13S,17R)-17-(furan-3-yl)-4,4,8,10,13-pentamethyl-3,16-dioxo-6,7,9,11,12,17-hexahydro-5H-cyclopenta[a]phenanthren-7-yl] acetate |
| Epoxyazadiradione | C<sub>28</sub>H<sub>34</sub>O<sub>6</sub> |  | (1S,2aR,3aR,3bS,4R,5aR,9aR,9bR,11aS)-1-(furan-3-yl)-3b,6,6,9a,11a-pentamethyl-2,7-dioxo-1,2,2a,3b,4,5,5a,6,7,9a,9b,10,11,11a-tetradecahydronaphtho[1',2':6,7]indeno[1,7a-b]oxiren-4-yl acetate |
| Azdirone | C<sub>28</sub>H<sub>36</sub>O<sub>4</sub> |  | [(5R,7R,8R,9R,10R,13S,17R)-17-(furan-3-yl)-4,4,8,10,13-pentamethyl-3-oxo-5,6,7,9,11,12,16,17-octahydrocyclopenta[a]phenanthren-7-yl] acetate |

**S1**: m/z: 450.52, C28H34O5.

1H nmr (400 MHz, CDCl3); δ(ppm): 7.48 (1H, m), 7.44 (1H, m), 7.14 (1H, d, J=10.18Hz), 6.28 (1H, m), 5.90 (1H, d, J=10.18 Hz), 5.89 (1H, s), 5.33 (1H, m), 3.43 (1H, s), 2.50 (1H, m), 2.21 (1H, m), 1.96 (3H, s), 1.35 (3H, s), 1.26 (3H, s), 1.11 (3H, s), 1.10 (3H, s), 1.04 (3H, s). 13C nmr (100 MHz, CDCl3); δ(ppm): 204.96, 203.96, 192.24, 169.58, 156.67, 142.80, 141.67, 125.96, 123.33, 118.44, 111, 73.92, 60.76, 47.97, 46.15, 44.09, 40.01, 38.23, 30.36, 27.00, 26.45, 26.30, 23.46, 21.28, 20.95, 19.02, 15.83.

**S2**: m/z: 484.52, C28H36O7.

1H NMR (400 MHz, CDCl3); δ(ppm ): 7.43 (d, J = 16.0 Hz, 2H), 6.26 (s, 1H), 5.85 (s, 1H), 5.34 – 5.21 (m, 1H), 4.70 (s, 1H), 3.96 (d, J = 19.0 Hz, 2H), 3.38 (s, 1H), 3.01 (d, J = 10.7 Hz, 1H), 2.55 (s, 1H), 2.27 (dd, J = 13.2, 2.3 Hz, 1H), 2.12 – 2.01 (m, 2H), 2.00 – 1.96 (m, 1H), 1.94 (s, 3H), 1.88 (d, J = 14.7 Hz, 1H), 1.82 – 1.66 (m, 2H), 1.33 (s, 3H), 1.29 (s, 3H), 1.12 (s, 3H), 1.08 (s, 3H), 1.03 (s, 3H). 13C NMR (100 MHz, CDCl3); δ(ppm ): δ 214.31, 205.53, 193.51, 169.99, 142.87, 141.75, 123.43, 118.74, 111.35, 74.22, 71.39, 60.70, 48.49, 46.73, 44.16, 42.28, 41.18, 35.55, 30.46, 26.80, 25.84, 24.08, 23.44, 21.16, 15.84, 15.35, 0.14.

**S3**: m/z: 524.52, C31H40O7.

^1^H NMR (400 MHz, CDCl3); δ(ppm ): δ 7.46 (s, 1H), 7.42 (s, 1H), 6.27 (s, 1H), 5.91 (d, *J* = 7.3 Hz, 1H), 5.25 (s, 2H), 4.39 (d, *J* = 7.6 Hz, 2H), 4.15 (d, *J* = 8.7 Hz, 2H), 3.40 (s, 1H), 3.04 (d, *J* = 13.7 Hz, 2H), 2.54 (d, *J* = 16.9 Hz, 2H), 1.96 (s, 5H), 1.87 (d, *J* = 8.9 Hz, 6H), 1.81 – 1.66 (m, 6H), 1.52 (s, 5H), 1.32 (s, 5H), 1.26 (d, *J* = 7.3 Hz, 8H), 1.21 (s, 8H), 1.07 (s, 9H), 0.80 (d, *J* = 18.6 Hz, 6H). ^13^C NMR (100 MHz, CDCl3); δ(ppm ): *δ* 212.16, 205.58, 193.19, 169.69, 142.77, 141.69, 123.42, 118.75, 111.31, 110.40, 82.00, 78.42, 74.46, 60.71, 48.25, 45.24, 43.39, 40.20, 38.25, 32.56, 30.49, 29.43, 26.07, 25.91, 25.13, 23.85, 21.64, 21.01, 15.47, 15.17, 0.07.

**S4**: 1) m/z: 408.52, C26H32O4

^1^H NMR (400 MHz, CDCl3); δ(ppm ): *δ* 7.45 (d, *J* = 15.5 Hz, 1H), 7.12 (d, *J* = 10.1 Hz, 1H), 6.27 (s, 1H), 6.03 (s, 1H), 5.87 (d, *J* = 10.5 Hz, 1H), 4.19 (s, 1H), 3.45 (s, 1H), 2.54 (q, *J* = 8.8 Hz, 1H), 2.48 (s, 1H), 2.12 – 1.95 (m, 4H), 1.85 (t, *J* = 12.0 Hz, 4H), 1.29 (s, 3H), 1.24 (d, *J* = 9.3 Hz, 8H), 1.17 (s, 5H), 1.11 (s, 5H), 1.03 (s, 3H). ^13^C NMR (101 MHz, CDCl3); δ(ppm ): *δ* 205.50, 204.76, 194.21, 157.30, 142.99, 141.80, 126.02, 123.56, 118.53, 111.30, 71.86, 61.01, 48.43, 46.73, 44.71, 44.37, 40.35, 36.67, 30.55, 29.85, 27.27, 26.83, 26.20, 25.50, 21.62, 19.26, 15.87, 0.15.

**S5:** m/z: 467.52, C28H32O6

1H nmr (400 MHz, CDCl3); δ(ppm): 7.56 (1H, m), 7.40 (1H, m), 7.17 (1H, d, J=10.04 Hz), 6.24 (1H, m), 5.88 (1H, d, J=10.04 Hz), 4.73 (1H, m), 3.90 (1H, s), 3.41 (1H, s), 2.63 (1H, m), 2.20- 2.15 (2H, m), 2.03 (3H, s), 1.22 (3H, s), 1.21 (3H, s), 1.08 (3H, s), 1.07 (3H, s), 1.04 (3H, s). 13C nmr (100 MHz, CDCl3); δ(ppm): 208.40, 204, 169, 157, 142.43, 141.53, 125.76, 116.51, 110, 73, 72, 57.18, 50.94, 46.67, 44.22, 43.12, 42.53, 39.69, 39.65, 29.09, 27.02, 24.82, 24.20, 21.26, 20.99, 19.82, 19.39, 16.06.

**S6:** m/z: 458.52, C28H42O5

1H NMR (400 MHz, Chloroform-d) δ 5.73 (s, 1H), 5.67 (s, 1H), 5.25 (s, 1H), 4.57 – 4.45 (m, 0H), 3.99 (d, J = 8.2 Hz, 1H), 3.86 – 3.78 (m, 1H), 3.74 (p, J = 5.8, 5.0 Hz, 2H), 3.65 (dt, J = 15.8, 7.6 Hz, 1H), 3.59 – 3.46 (m, 1H), 2.60 (d, J = 17.1 Hz, 2H), 2.49 – 2.38 (m, 2H), 2.34 (s, 1H), 2.26 – 2.17 (m, 2H), 2.17 – 2.05 (m, 4H), 1.95 (s, 9H), 1.90 – 1.74 (m, 9H), 1.34 (d, J = 9.5 Hz, 5H), 1.27 (d, J = 12.0 Hz, 15H), 1.11 (s, 5H), 1.04 (d, J = 11.7 Hz, 10H).

**S7:** m/z: 456.52, C28H40O5

^1^H NMR (400 MHz, CDCl3); δ(ppm ): δ 7.32 (d, *J* = 8.5 Hz, 2H), 7.17 (s, 2H), 6.67 (s, 1H), 6.49 (d, *J* = 10.4 Hz, 1H), 6.36 (s, 1H), 6.11 (d, *J* = 15.3 Hz, 1H), 5.86 (s, 1H), 5.31 (d, *J* = 13.1 Hz, 2H), 3.97 (s, 1H), 3.54 (s, 1H), 2.27 (ddt, *J* = 31.4, 21.5, 12.0 Hz, 2H), 2.01 – 1.87 (m, 11H), 1.64 (dd, *J* = 19.7, 10.8 Hz, 7H), 1.36 (s, 4H), 1.32 (s, 3H), 1.24 (d, *J* = 9.4 Hz, 8H), 1.20 (s, 2H), 1.15 (d, *J* = 9.2 Hz, 10H), 1.08 (s, 6H), 0.89 (d, *J* = 7.2 Hz, 2H).

**S8:** m/z: 456.52, C28H40O5

^1^H NMR (400 MHz, CDCl3); δ(ppm ): δ 5.75 (s, 1H), 5.26 (s, 1H), 4.10 – 3.99 (m, 1H), 3.86 (dd, *J* = 8.7, 3.7 Hz, 1H), 3.83 – 3.74 (m, 1H), 3.39 – 3.30 (m, 1H), 2.76 – 2.55 (m, 2H), 2.50 – 2.39 (m, 1H), 2.34 (q, *J* = 8.5 Hz, 1H), 2.30 – 2.21 (m, 1H), 2.04 (d, *J* = 10.3 Hz, 1H), 1.94 (s, 5H), 1.85 (d, *J* = 9.2 Hz, 3H), 1.50 (s, 1H), 1.26 (d, *J* = 13.1 Hz, 17H), 1.09 (s, 3H), 1.03 (d, *J* = 11.1 Hz, 7H). ^13^C NMR (100 MHz, CDCl3); δ(ppm ): *δ* 216.16, 207.27, 192.27, 169.87, 132.29, 132.19, 128.72, 128.60, 123.61, 77.48, 77.37, 77.16, 76.84, 74.52, 71.83, 68.15, 66.21, 48.32, 47.64, 47.04, 43.86, 42.36, 38.61, 37.57, 33.96, 32.07, 31.58, 31.54, 31.08, 30.32, 29.84, 29.80, 29.51, 28.48, 26.01, 25.91, 25.19, 24.05, 22.84, 21.21, 21.16, 15.93, 15.13, 14.28, 1.17, 0.14.

**S9:** m/z: 521.52, C29H37O6

1H NMR (400 MHz, CDCl3); δ(ppm ): δ 7.51 (s, 1H), 7.12 (d, J = 10.0 Hz, 1H), 6.43 (s, 1H), 5.90 (s, 2H), 5.33 (s, 1H), 4.54 (s, 1H), 2.50 (s, 3H), 2.48 (s, 1H), 2.22 (s, 1H), 2.19 (s, 1H), 2.08 (s, 2H), 2.01 (s, 2H), 1.97 (s, 4H), 1.92 (d, J = 18.7 Hz, 3H), 1.37 (s, 5H), 1.25 (s, 30H), 1.08 (d, J = 12.1 Hz, 11H), 0.88 (s, 4H). ^13^C NMR (10 MHz, CDCl3); δ(ppm ): δ 205.36, 204.22, 193.63, 189.46, 169.72, 157.05, 150.42, 144.47, 127.74, 125.97, 123.40, 115.82, 74.19, 59.95, 50.21, 46.35, 45.02, 44.24, 40.17, 38.09, 32.08, 30.55, 29.83, 29.51, 27.35, 27.09, 26.63, 23.59, 22.84, 21.45, 21.16, 19.17, 15.96, 14.27.

**S10**: m /z: 436.52, C28H40O5

1H NMR (400 MHz, CDCl3); δ(ppm ): δ 1.15 (3H, s), 1.18 (3H, s), 1.26 (3H, s), 1.29-1.51 (8H, 1.40 (dddd, J = 13.1, 10.3, 9.9, 3.0 Hz), 1.30 (s), 1.44 (ddd, J = 13.1, 10.3, 2.9 Hz), 1.37 (s)), 1.63-1.75 (2H, 1.69 (dddd, J = 13.1, 2.9, 2.8, 1.8 Hz), 1.70 (ddd, J = 13.1, 3.0, 2.8 Hz)), 1.77- 1.97 (3H, 1.91 (ddd, J = 13.1, 8.1, 3.3 Hz), 1.81 (dd, J = 10.1, 3.3 Hz), 1.87 (dd, J = 9.9, 1.8 Hz)), 2.05-2.33 (5H, 2.12 (ddd, J = 13.1, 10.1, 3.1 Hz), 2.26 (ddd, J = 13.6, 7.9, 4.6 Hz), 2.11 (s)), 2.35 (1H, ddd, J = 13.6, 5.3, 4.3 Hz), 3.55 (1H, dd, J = 7.9, 5.3 Hz), 5.87-5.96 (2H, 5.93 (dd, J = 8.1, 3.1 Hz), 5.89 (dd, J = 4.6, 4.3 Hz)), 6.27 (1H, dd, J = 1.8, 1.1 Hz), 6.53 (1H, d, J = 9.5 Hz), 7.00-7.07 (2H, 7.04 (d, J = 9.5 Hz), 7.01 (dd, J = 1.1, 0.9 Hz)), 7.30 (1H, dd, J = 1.8, 0.9 Hz). ^13^C NMR (10 MHz, CDCl3); δ(ppm ): 203, 170, 159, 155,140, 124, 39.4, 20.9, 23.54, 32.05, 37.3, 34.3, 73.8, 111.4, ,23.5, 17.71, 47.4, 43.35, 119.55, 51.5, 142.63, 23.542, 24.9, 44.07, 20.95

**S11:** m/z: 482.52, C28H34O7

1H NMR (500 MHz, CDCl3); δ(ppm ): δ 7.40 (d, J = 1.7 Hz, 2H), 7.09 (d, J = 10.2 Hz, 1H), 6.41 – 6.29 (m, 1H), 5.85 (d, J = 10.3 Hz, 1H), 5.60 (s, 1H), 4.54 (dd, J = 3.8, 2.1 Hz, 1H), 3.51 (s, 1H), 2.47 (dd, J = 12.8, 6.1 Hz, 1H), 2.15 (dd, J = 13.3, 2.5 Hz, 1H), 2.09 (s, 3H), 1.99 (ddq, J = 10.2, 7.2, 3.4 Hz, 1H), 1.93 (dt, J = 15.0, 3.2 Hz, 1H), 1.88 – 1.78 (m, 2H), 1.77 – 1.69 (m, 1H), 1.58 (ddd, J = 13.5, 10.6, 2.5 Hz, 1H), 1.23 (s, 3H), 1.21 (s, 3H), 1.14 (s, 3H), 1.05 (d, J = 3.5 Hz, 6H). 13C NMR (126 MHz, CDCl3); δ(ppm ): δ 204.09, 169.99, 167.57, 157.11, 143.19, 141.29, 126.08, 120.52, 109.97, 78.38, 73.33, 69.90, 56.98, 46.13, 44.15, 42.73, 40.14, 39.62, 38.83, 27.27, 26.10, 23.37, 21.30, 21.19, 19.86, 18.42, 17.84, 15.08.

We use S1 as Azadiradione (AZD), and S5 is Epoxy-azadiradione.

### 3.2 Epoxy-azadiradione activates cellular HSF1 with higher efficacy than Azadiradione and ameliorates protein aggregation-induced toxicity in cells

To understand the functional derivative(s) of AZD, we analyzed the cellular HSP70A1A-inducing activity of the compounds (Figure 1) using an RT-qPCR assay. Levels of HSP70A1A were also estimated in cells treated with mock as a negative control and heat shock as a positive control. As shown in Figure 2A, compound S5, an epoxy derivative of AZD, showed two-fold more activity than AZD. Another compound in this collection of derivatives, compound S11, showed activity comparable to that of AZD. A higher Epoxy activity compared to AZD was observed, as indicated by higher HSP70 protein levels (the product of the HSP70A1A gene) in those cells (Figure 2C). We have chosen to further analyze the activity of the Epoxy derivative.

**FIGURE: 2.**
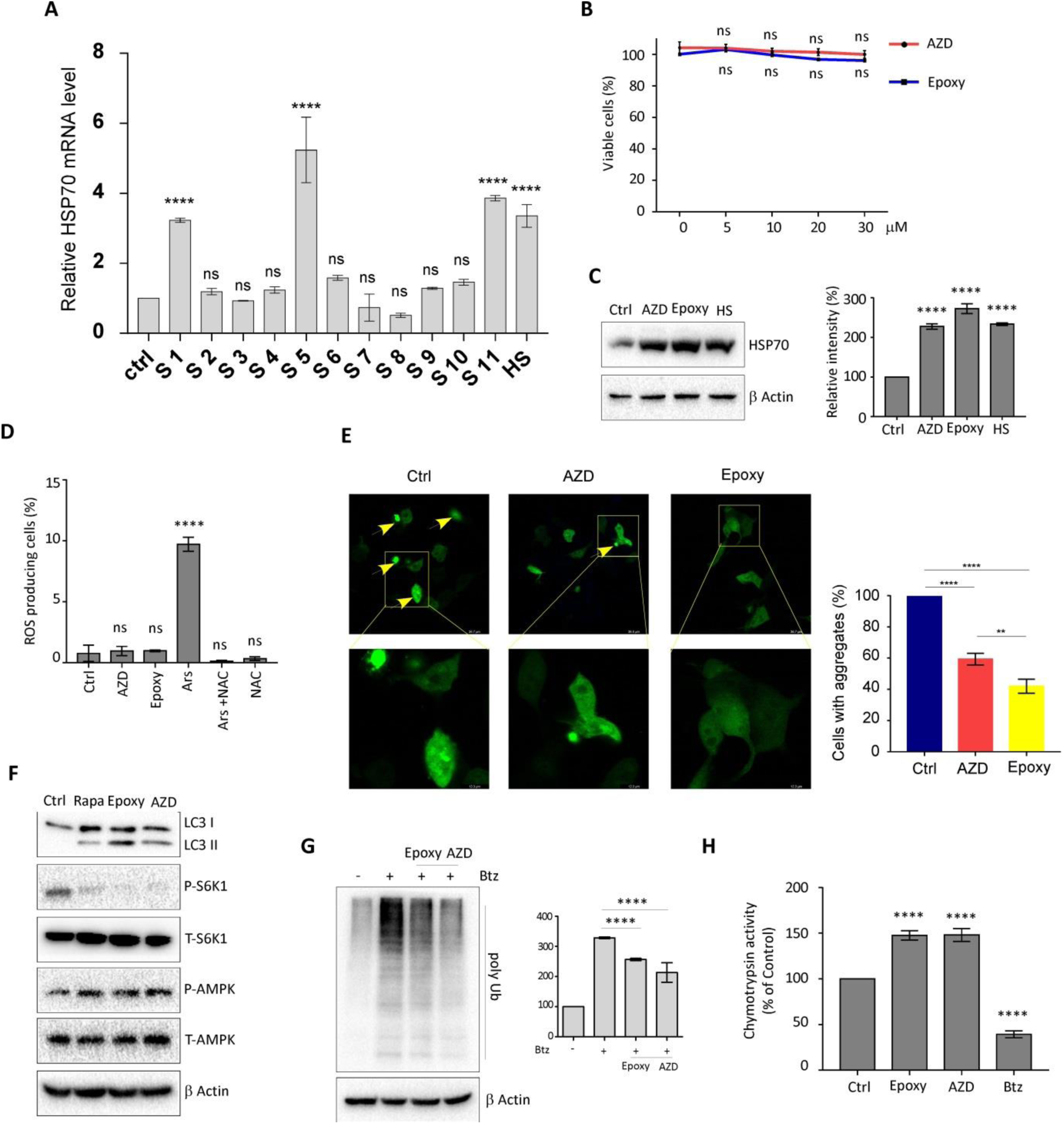
**Derivatives of azadiradione exhibited differential activities in HEK293 cells.** [A] Relative HSP70A1A inducing activities of azadiradione estimated by RT-qPCR assays as described in the materials and methods. Cells were treated with 20 µM of compounds for 2 h, or subjected to heat shock (HS) for 30 min prior to processing for RT-qPCR, as described in the Materials and Methods section. [B] Comparison of the effect of epoxy-azadiradione (Epoxy) with azadiradione (AZD) on cellular viability. Cells were treated with the indicated doses of the compounds before being processed for MTT assays. [C] Representative immunoblots showing the HSP70-inducing activities of azadiradione (AZD), Epoxy-azadiradione (Epoxy), and heat shock (HS). The bar graph on the right represents the densitometric quantification of the bands. Cells were treated with the indicated concentrations of the compounds overnight or for 30 minutes with HS, followed by a 6-hour recovery period, and then harvested for protein analysis. β-Actin level was determined as an internal loading control (Ctrl). [D] Azadiradione (AZD) or epoxy-azadiradione (Epoxy) does not induce ROS levels in cells as estimated by the DCFDA method as described in the Materials and Methods. Representative bar graph indicating the ROS levels in cells pretreated as indicated. Cells were treated overnight with the compounds or for 2 h with 20 µM arsenite (Ars) as a positive control before processing for the assay. Cells were pretreated with N-acetylcysteine (NAC) for 2 h alone or before Ars treatment. [E] Effect of azadiradione (AZD) and epoxy-azadiradione (Epoxy) on the dissolution of aggregates due to overexpression of ataxin-130Q as observed by GFP fluorescence level analysis. Cells were treated with the vehicle (Ctrl), 20 µM AZD, or Epoxy for 24 h before being processed for fluorescence imaging. The bar graph on the right shows the estimated assay result.[F] Representative immunoblots showing the relative effect on mTORC1 of its inhibitor rapamycin, with azadiradione (AZD) and epoxy-azadiradione (Epoxy) on the expression level of LC3II/I, phospho-S6K (p-S6K), and phospho-AMPK (p-AMPK). Cells were pretreated with 500 nM rapamycin for 24 h or 20 µM of AZD or Epoxy, and protein extracts were analyzed by western blot using antibodies against LC3, p-S6K, and p-AMPK. Treatment with Epoxy and AZD increased LC3II levels and decreased p-S6K levels compared with rapamycin, indicating enhanced autophagy and further suppression of mTORC1 signalling. Β Actin was used as a loading control. [G] Representative immunoblot showing the level of poly-ubiquitinated proteins in cells pretreated with azadiradione (AZD) and epoxy-azadiradione (Epoxy). Cells were treated with vehicle (Ctrl), 20 µM AZD, 20 µM Epoxy, or bortezomib (Btz) for 24 h. Cells were pre-treated with Bortezomib (Btz 100 nM) for 2 hours. The bar graph on the right shows the densitometric quantification of the bands. [H] Enzymatic quantitation of chymotrypsin-like activity (20s proteasome activity) in cells treated with azadiradione (AZD) and epoxy-azadiradione (Epoxy). Cells were treated with vehicle (Ctrl), 20 µM AZD, or Epoxy for 24 h, or bortezomib (Btz) for 2 h as a positive control, before being processed for the assay. All Data are presented as mean ± SD of three independent experiments. P value with **** P<0.0001 respectively

The sensitivity of human embryonic kidney (HEK293) and mouse neuroblastoma (Neuro-2A) cells to different concentrations of AZD and Epoxy was tested by MTT assay. Results show that the viability of HEK293 cells was not compromised by up to 30 μM of either compound in a 24-hour incubation period (Figure 2B). We then tested whether treating cells with AZD and Epoxy induces cellular reactive oxygen species (ROS) levels. This test was conducted because many plant-derived HSF1 activators, such as celastrol, sulforaphane, and gedunin, have been shown to work by inducing cellular ROS (Westerheide and Morimoto 2005; Gan et al. 2010; Patwardhan et al. 2013; Nelson et al. 2016). Estimation was carried out using the DCFDA method, which showed minimal or no elevation of ROS production in AZD- or Epoxy-treated cells (Figure 2D). Next, we tested Epoxy for its ability to reduce the aggregation levels of an aggregation-prone ataxin130QGFP fusion protein transiently overexpressed in Neuro-2A cells compared to AZD-and vehicle-treated cells. Cells overexpressing the fusion protein were treated overnight with 20 µM of either AZD or Epoxy. Results indicated a gradual decrease in ataxin130Q-GFP aggregate levels in Neuro-2A cells treated with both AZD and Epoxy, as measured by GFP fluorescence levels (Figure 2E). Notably, the Epoxy action in reducing ataxin130Q-GFP aggregate levels was more effective than that of AZD (Figure 2E). Densitometric analyses showed that the induction of HSP70A1A transcript levels in Epoxy- and AZD-treated HEK293 cells was approximately 5-fold and 2.5-fold, respectively, in agreement with the reduction in ataxin130Q-GFP aggregation in Epoxy-treated samples, indicating a correlation between Epoxy-mediated protein aggregation-resolving activity and the elevation of HSP70 expression levels (Figure 2C).

### 3.3 Epoxy-azadiradione induces autophagy by activation of AMPK and inhibits mTORC1

Autophagy is a critical mechanism for clearing cellular protein aggregates (Menzies et al., 2017). To test whether Epoxy and AZD induce the cellular autophagy process, Neuro-2A cells were treated with an equivalent concentration of the compounds, using Rapamycin, an inhibitor of mTORC1, as a positive control.

The effect of the treatment on the cellular level of AMPK, a critical modulator of mTORC1 function, was also tested during this process (Kim et al., 2011; Menzies et al., 2017). Immunoblot experiments showed that both Epoxy and AZD treatments significantly induced cellular autophagy as suggested by upregulation of LC3II cleavage and inhibition of S6K1 phosphorylation as a downstream target of mTORC1. Upregulation of AMPK by Epoxy, like AZD, suggested its involvement in mTORC1 inhibition in both conditions in the treated cells (Figure 2F).

### 3.4 Epoxy-azadiradione activates cellular protein aggregation clearance mechanisms

We measured the chymotrypsin activity of the 20S proteasome in the lysates of cells pre-treated with either vehicle or bortezomib, a proteasome inhibitor, plus Epoxy or AZD (Kisselev et al. 2006). As revealed, Epoxy could induce the chymotrypsin activity of the 20S proteasome. In the ubiquitin-proteasome system, polyubiquitinated proteins are degraded by the 20S proteasome. We next investigated the involvement of the proteasome pathway in AZD- and Epoxy-treated cells by examining the abundance of polyubiquitinated (polyUb) proteins and the enzymatic activity of the proteasome. As revealed in the immunoblots, both the signatures of the ubiquitin-proteasome pathway were stimulated in the cells treated with the compounds (Figure 2G & H).

### 3.5 Epoxy-azadiradione ameliorates PD symptoms and pathology in MPTP-induced mice

Aggregation of α-synuclein in the substantia nigra pars compacta (SNPc), and loss of dopaminergic neurons (DA) are pathological hallmarks in Parkinson’s disease (PD) (Mehra et al. 2019). The loss of function of chaperone proteins and upregulation of chaperone activity were directly correlated with the loss and restoration of dopaminergic neurons in PD patients. Our results with the cell culture model, indicating efficient Epoxy-induced activation of cellular HSF1 and HSP70, have encouraged us to test the effect of Epoxy in a mouse model of PD developed by treatment with MPTP. Swiss albino mice pre-treated with MPTP (10 mg/kg) were treated with Epoxy on alternate days for 20 days. Separate groups of mice were also treated with the same dose of AZD as a control. The development of PD and the efficacy of Epoxy and AZD on reversing PD symptoms were assessed using the grip test and body weight test (Figure 3). Mouse brains were collected at the end of the treatments, and the levels of Hsp70, α-synuclein, and tyrosine hydroxylase (TH) in SNPc tissues from treated and control samples were determined by immunoblot experiments. MPTP-treated mice, after treatment with Epoxy, exhibited significantly reduced PD symptoms, as assessed by the animals’ grip strength and body weight (Figure 3A-C). TH, a hallmark of dopaminergic neurons/ SNPc and a key enzyme in the dopamine biosynthesis pathway, is progressively lost in PD animals. As expected, Epoxy treatment significantly restored TH levels in SNPc in PD animals compared with AZD- or vehicle-treated mice, as assessed by immunohistochemistry (Figure 3D). Immunoblots performed with mouse brain samples showed upregulation of TH and Hsp70 levels and downregulation of α-synuclein levels (Figure 3E).

**FIGURE: 3.**
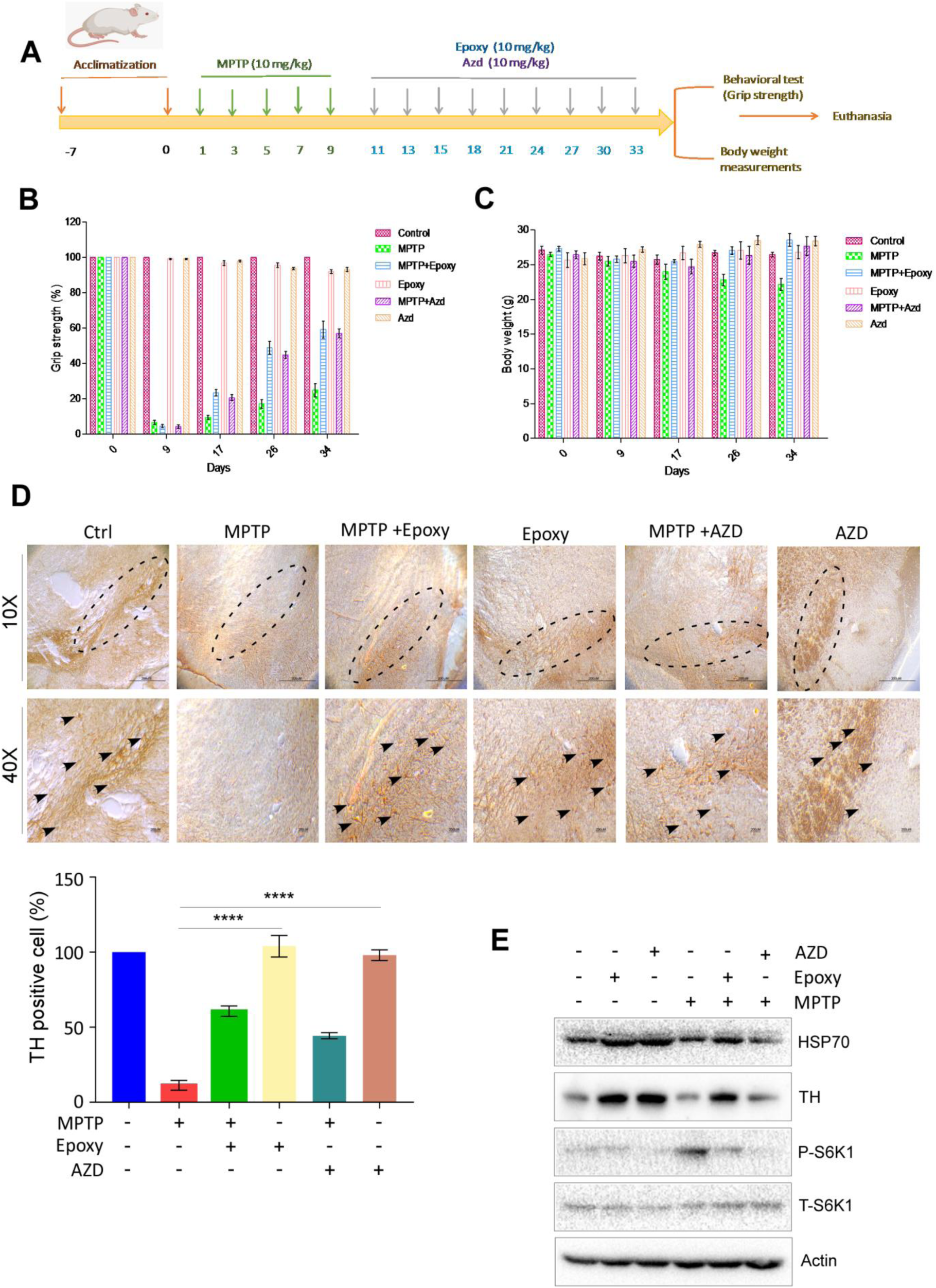
**Epoxyazadiradione ameliorated Parkinson’s disease developed in Swiss albino mice.** [A] Schematic representation of the workflow of the experiment for testing the effect of azadiradione (AZD) and epoxy-azadiradione (Epoxy) on Parkinson’s disease in Swiss albino mice.[B-C] Representative bar graph showing the effect of treatment with the vehicle (Ctrl), Epoxy-azadiradione (Epoxy), or Azadiradione (AZD) on the recovery of lost gripping ability and body weight in MPTP-treated mice. [D] Representative images of brains isolated from mice pretreated with the vehicle (Ctrl), Epoxy-azadiradione (Epoxy), or Azadiradione (AZD), stained with tyrosine hydroxylase (TH) antibody. TH levels in the substantia nigra pars compacta in the sections are marked by broken ovals and arrowheads, indicating the dopaminergic neurons. [E] Representative immunoblots showing the levels of indicated factors in the lysates of brain isolated from animals pretreated as described in panel A and Materials and Methods. Densitometric quantitation of bands is shown as a bar graph. β-Actin level was determined as an internal loading control. All Data are presented as mean ± SD of three independent experiments. P value with **** P<0.0001, respectively.

### 3.6 Epoxy-azadiradione induced DNA binding affinity of HSF1

We previously demonstrated that AZD enhances the DNA binding affinity of HSF1 (Nelson et al., 2016). To better understand the differential activities of AZD and Epoxy on the transcriptional activity of HSF1 (Figure 2A), we have measured the Kd of binding of the full-length HSF1 to the heat shock element (HSE) in the presence of AZD or Epoxy by fluorescence polarization (FP) assays in vitro. We also included other structural derivatives (Figure 1) that were inactive in inducing HSF1 transcription function to assess whether stimulation of HSF1-HSE interaction correlates with their transcriptional functions (Figure 1). As shown in Figure 4, Epoxy induced HSF1-HSE interaction by 2-fold higher affinity than that of AZD (3.4590 nM vs 8.8090 nM). As expected, the other derivatives did not show any net stimulation of HSF1-HSE interaction, consistent with their negligible intracellular transcriptional activities (Figure 2A).

**FIGURE: 4.**
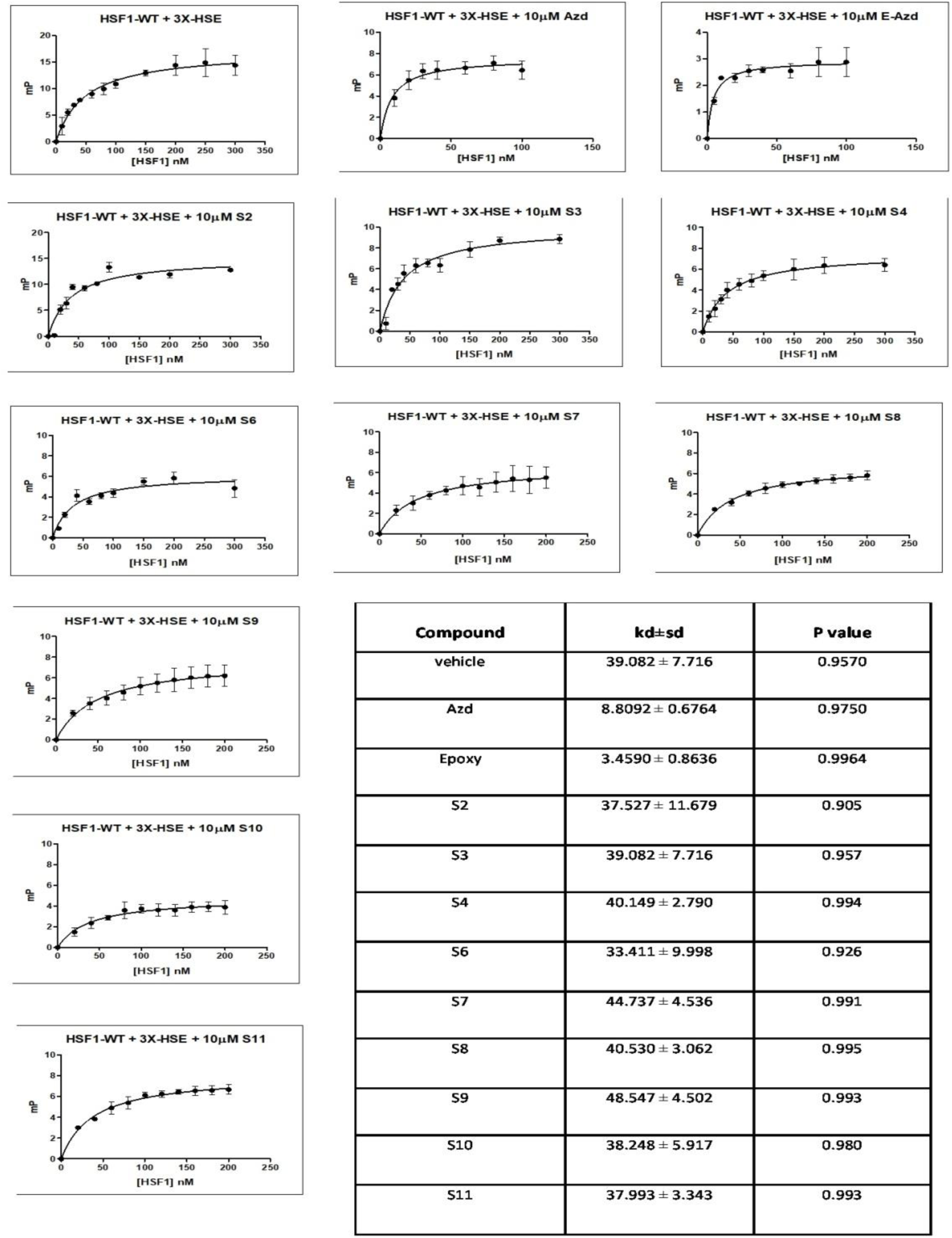
**Epoxy-azadiradione induces higher affinity binding of HSF1 to its target DNA element compared to AZD.** The affinities of full-length HSF1 protein to its recognition sequence, heat shock element (HSE, as a double-stranded DNA oligonucleotide at 10 nM), in the presence of AZD and its various structural derivatives were estimated by fluorescence polarization techniques. AZD and its structural derivatives were titrated in the reaction with millipolarization values taken at various HSF1 concentrations. The KD values were calculated by plotting curves from three independent experiments and performing a one-site specific binding fit in GraphPad Prism software. Epoxy demonstrated a lower nanomolar affinity (kd) with HSF1 protein than other derivatives. The table represents the collective data of the kd of AZD, Epoxy, and its other structural derivatives as indicated.

### 3.7 Molecular docking suggested the assembly of more stable HSF1-HSE complexes by Epoxy-azadiradione compared to Azadiradione

The interactions of AZD and Epoxy with the DNA binding domain of HSF1 (HSF1-HSE) structure, available in the Protein Data Bank (PDB), along with their binding affinities, were estimated by molecular docking to understand the potential molecular bases of their effects in enhancing cellular HSF1 function (Neudegger et al., 2016). Energetically preferred 3D conformers of AZD and Epoxy were used in the docking analyses described in the Materials and Methods to generate docking solutions (Figure 5) to pool together the best-ranked docked solutions (top 15) for further scoring using the AutoDock Vina program (Trott and Olson et al., 2010). Docked complexes having RMSD <= 2.5Å are grouped into a single cluster. Snapshot docking scores of AZD conformers were found to be higher at both probable binding sites compared to those of Epoxy (Figure 5A). However, further molecular dynamics-based energy estimation analysis revealed that although AZD possessed higher initial binding/docking scores compared to Epoxy, the energetic stability of the Epoxy complexes is higher as indicated by much higher decrease of binding free energy (ΔG) values estimated from an ensemble of intermediate docked complex structures (Fig. 5B). Intermediate complex structures extracted at midway (5 ns) and end (10 ns) of the simulation runs were utilized to identify probable interacting HSF1 and DNA residues with the respective ligands/activator. Figures 5 and 6 provide pictorial and tabular views of the probable interacting residues.

**FIGURE: 5.**
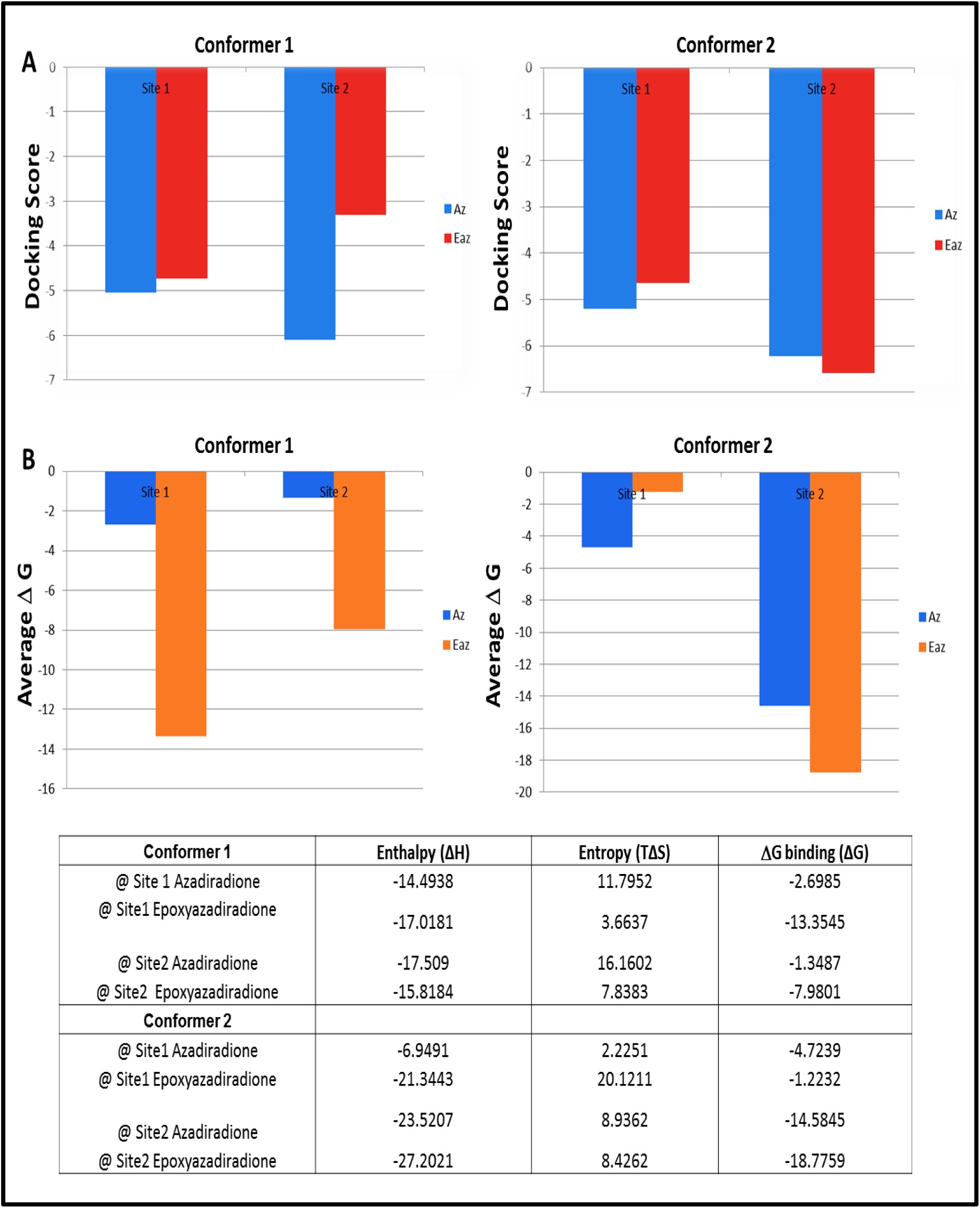
**Comparison of docking scores and free energies of the azadiradione (AZD) and epoxy-azadiradione (Epoxy) complexes.** Docking scores of the AZD and Epoxy complexes derived by the meta-docking approach were compared in panel A, while the average free energies (ΔG) of the ensemble docked complexes are compared in panel B. Entropy and enthalpy components of the binding free energies (ΔG) are provided in the lower panel.

**FIGURE: 6.**
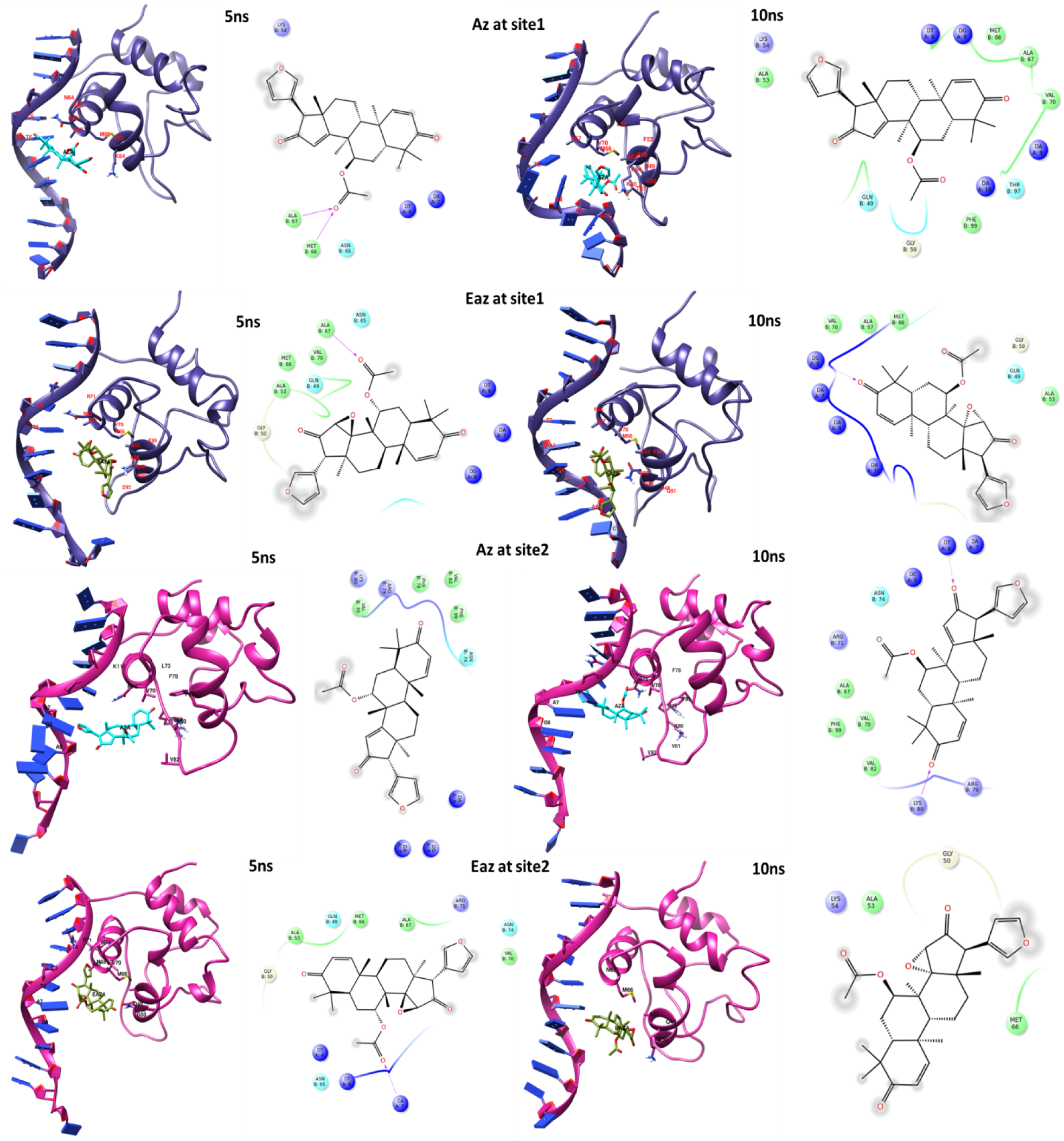
**Probable interactive residues with respect to the docked azadiradione (AZD) and epoxy-azadiradione (Epoxy).** Probable binding residues with respect to docked azadiradione (AZD) and epoxy-azadiradione (Epoxy) are shown in cartoon and LIGPLOT views within the docked intermediate complex structures extracted at 5 and 10 ns of the simulation, respectively.

## 4 Discussion

Various naturally occurring small molecules from Azadirachta indica have been reported to have neuroprotective activity (Sandhir et al. 2021). The neuroprotective action of Epoxy was studied in Alzheimer’s disease (Gorantla et al. 2020). Several limonoids, including Gedunin, Epoxy, and AZD, were tested in vitro and in silico for their ability to inhibit tau aggregation and promote the dissolution of tau fibrils. Epoxy was the most neuroprotective basic limonoid, disintegrating tau fibrils (63.5%) and activating HSF1 protein (Gorantla et al., 2020). The findings suggest that Epoxy carries a potential therapeutic prospect for neurodegenerative diseases involving tauopathies as well. Several earlier reports showed Epoxy as an anticancer (Kumar et al., 2018; Lakshmi et al., 2021; Rai et al., 2020; Shilpa et al., 2017), anti-inflammatory (Alam et al., 2012), anti-plasmodial (Ashok Yadav et al., 2017), and antifungal molecule (Govindachari et al., 1998). Epoxy demonstrated anticancer potential by suppressing breast tumour growth in triple-negative breast cancer (TNBC) and ER+ cells, inducing apoptosis, and inhibiting PI3K/AKT-mediated mitochondrial potential both in vitro and in vivo (Kumar et al., 2018). Another report suggests that Epoxy plays a role in TNBC by inducing G2/M phase arrest in the cell cycle, inducing anoikis, and inhibiting NF-kB nuclear translocation (Lakshmi et al., 2021). Epoxy was previously reported to inhibit cervical cancer (Shilpa et al., 2017) and head and neck squamous cell carcinoma by blocking the translocation of NF-κB (Rai et al., 2020).

Our objective in this endeavour is to investigate the structure-activity or pharmacophore of AZD through the modification of functional groups, thereby defining the structural requirements for activating the heat shock-like response. Through analysing the Hsp70-inducing activity of different structural derivatives of AZD, we identified Epoxy as another compound with therapeutic potential for PD. Epoxy activity in cell and animal models appeared to be better than AZD. Our in vitro data suggested that Epoxy is at least 2-fold more efficient than AZD (Figures 1A, 4). The cellular tolerance of Epoxy is the same as that of AZD, suggesting that Epoxy is not more toxic than AZD (Figure 5). Docking scores indicated that the overall Epoxy binding score was very similar to the AZD binding energy. A 10 ns GROMACS molecular dynamics simulation, followed by gmx_MMPBSA-based binding energy estimation, shows that the Epoxy complex is energetically and entropically more stable compared to the AZD complex. The Epoxy group in Epoxy-AZD likely contributes to the complex’s stability (Figure S1). The basis of the higher efficacy of Epoxy is not yet clear. It is also interesting to investigate the molecular basis of why, among the ten derivatives, only AZD and Epoxy could activate HSF1 function.

In contrast, others that carry a structural alteration in other parts of the molecule lose activity, suggesting that these interactions with the HSF1-HSE complex are very stringent (Figure 4). Epoxidation of the double bond between positions 14 and 15 in AZD increased its efficacy (Figure 1)., whereas the saturation of double bonds diminished the effectiveness of AZD derivative (Figure 4). The saturation of the oxo group at position 16 inactivated the observed function (Azadirone, S11 Figure 1). We also observed that the alteration or acetylation of the carboxyl group inactivated the HSF1-inducing activity of AZD (data not shown).

Dopamine supplementation is the mainstay of current PD treatment. Although molecular chaperone upregulation through HSF1 functional activation is considered a potential therapy for this disease, the proinflammatory and pro-oncogenic nature of HSF1 has somewhat dampened enthusiasm. Nevertheless, AZD was demonstrated to inhibit cellular AKT activity, albeit through an HSF1-independent mechanism, indicating that AZD does not support the cellular proliferative pathway. It remains to be tested if Epoxy, like AZD, shows anti-AKT activity (Sareng, Dutta et al. 2025). Altogether, the anti-inflammatory and anticancer properties of Epoxy, as observed by other groups, may make it a safer molecule for PD therapy than other small-molecule HSF1 activators, many of which work through inducing cellular oxidative imbalance (Kumar et al., 2018; Lakshmi et al., 2021; Rai et al., 2020; Shilpa et al., 2017). Furthermore, many studies indicated an inverse relationship between various cancers and neurodegenerative disorders such as Parkinson’s disease (PD) and Alzheimer’s disease (AD) (Rojas et al. 2020). In other words, individuals with PD or AD have a comparatively lower risk of developing certain types of cancers (Inzelberg and Jankovic 2007).

AZD or Epoxy showed promising therapeutic potential for neurodegenerative diseases. A logical follow-up of this study could be to test the efficacy of Epoxy along with AZD in a small number of Parkinson’s patients. However, the compound’s limited abundance in neem seeds (∼100 mg/kg seed powder) has limited the scope of this option. Therefore, a logical strategy is to obtain and solve the cocrystal structure of AZD and Epoxy (and some inactive derivatives of AZD) with the HSF1-HSE complex, and to understand the interactions between the molecules responsible for their activities. A follow-up experiment to this study would be to collaborate with synthetic organic chemists and bioinformaticians to identify small molecules that recapitulate the interactions and to test their in vitro and in vivo activities, with the aim of considering them for clinical trials.

## Author contributions

S. C., N.D., H. R. S., N. G., A. M., I, M. K., data curation; N. D., H. R. S., S. C., I. M.K., formal analysis; N. D., H. R. S., R.K.G., J, M., Saikat, C., M. P., validation; N. D., H. R. S., and M. P. investigation; S. C., N. D., H. R. S., and M. P., visualization; N. D,, H. R. S., and S. C., methodology; S. C., N. D., and H. R. S., writing-original draft; M.P., R. K. G., J. M., Saikat C., resources; M. P. conceptualization; M. P. , R. G., J. M., Saikat, C., supervision; M. P., project administration; and S. C., N. D., I. M. K., writing the original draft, M. P., writing-review and editing.

## Acknowledgments

We thank the DBT, SERB, NIPER-K, and the Bose Institute for funding, as well as the faculty members of the Division of Molecular Medicine, now the Department of Biological Sciences at Bose Institute, for their support in carrying out this work. We gratefully acknowledge the help provided by Dr. Kuladip Jana during the animal experiment.

---

**FIGURE: S1.**
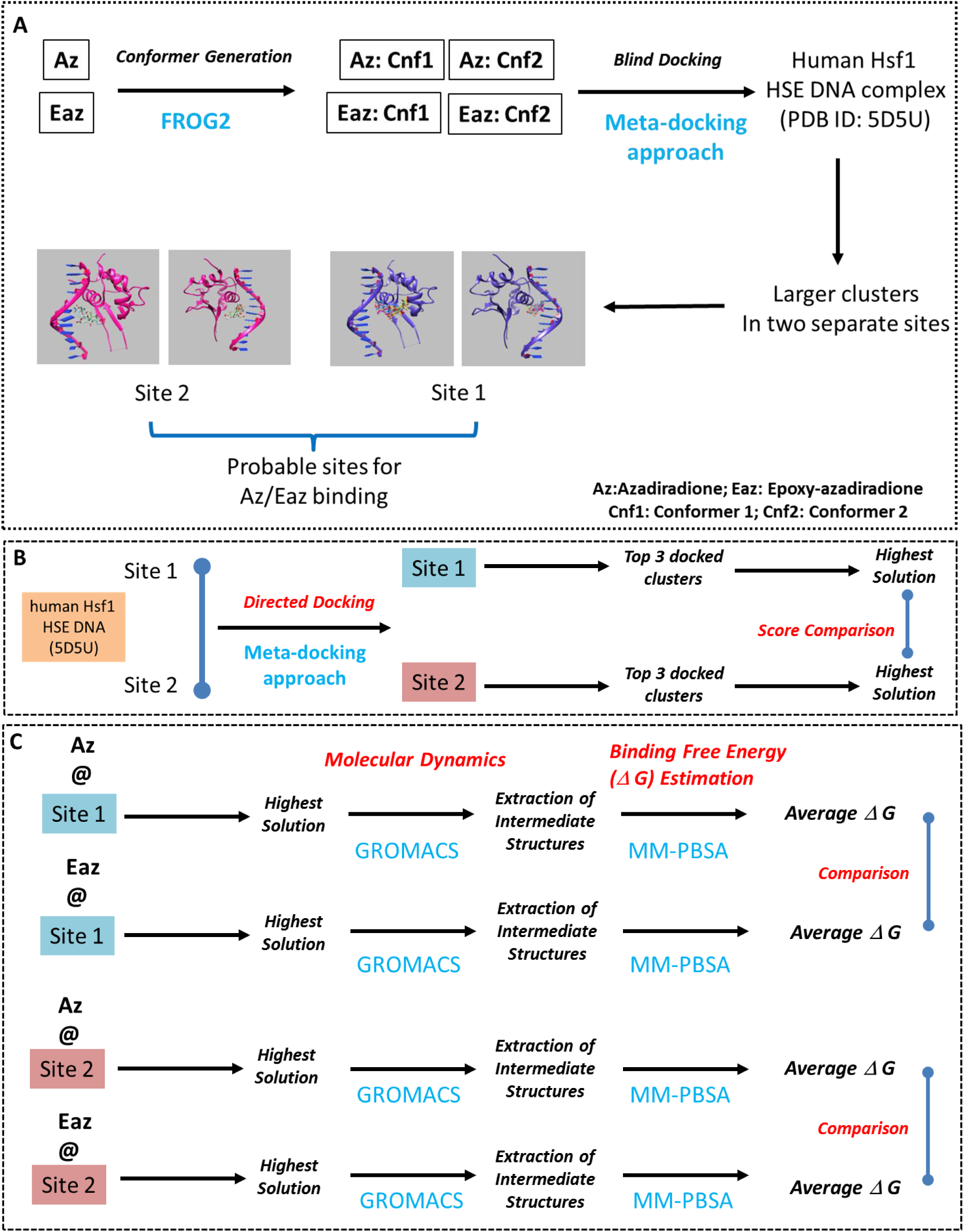

